# Genetic mapping and identification of a QTL determining tolerance to freezing stress in *Fragaria vesca* L.

**DOI:** 10.1101/2021.02.22.432243

**Authors:** Jahn Davik, Robert C. Wilson, Relindis G. Njah, Paul E. Grini, Stephen K. Randall, Muath K. Alsheik, Daniel James Sargent

**Affiliations:** Division of Biotechnology and Plant Health, Norwegian Institute of Bioeconomy Research, N-1431 Ås, Norway; Department of Biotechnology, Faculty of Applied Ecology, Agricultural Sciences & Biotechnology, Inland Norway University of Applied Sciences, NO-2318 Hamar, Norway; Section for Genetics and Evolutionary Biology (EVOGENE), Department of Biosciences, University of Oslo, NO-0316 Oslo, Norway; Department of Biology, Indiana University Purdue University Indianapolis, Indianapolis, IN, 46202, USA; Graminor Breeding Ltd., N-2322 Ridabu, Norway; Department of Genetics, Genomics and Breeding, NIAB-EMR, East Malling, Kent, ME19 6BJ, UK; Natural Resources Institute, University of Greenwich, Medway Campus, Central Avenue, Chatham Maritime, Kent, ME4 4TB, UK

## Abstract

Extreme cold and frost cause significant stress to plants which can potentially be lethal. Low temperature freezing stress can cause significant and irreversible damage to plant cells and can induce physiological and metabolic changes that impact on growth and development. Low temperatures cause physiological responses including winter dormancy and autumn cold hardening in strawberry (*Fragaria*) species, and some diploid *F. vesca* accessions have been shown to have adapted to low-temperature stresses. To study the genetics of freezing tolerance, a *F. vesca* mapping population of 142 seedlings segregating for differential responses to freezing stress was raised. The progeny was mapped using ‘Genotyping-by-Sequencing’ and a linkage map of 2,918 markers at 851 loci was resolved. The mapping population was phenotyped for freezing tolerance response under controlled and replicated laboratory conditions and subsequent quantitative trait loci analysis using interval mapping revealed a single significant quantitative trait locus on *Fvb2* in the physical interval 10.6 Mb and 15.73 Mb on the *F. vesca* v4.0 genome sequence. This physical interval contained 896 predicted genes, several of which had putative roles associated with tolerance to abiotic stresses including freezing. Differential expression analysis of the 896 QTL-associated gene predictions in the leaves and crowns from ‘Alta’ and ‘NCGR1363’ parental genotypes revealed genotype-specific changes in transcript accumulation in response to low temperature treatment as well as expression differences between genotypes prior to treatment for many of the genes. The putative roles, and significant interparental differential expression levels of several of the genes reported here identified them as good candidates for the control of the effects of freezing tolerance at the QTL identified in this investigation and the possible role of these candidate genes in response to freezing stress is discussed.

## Introduction

Climate change has resulted in greater instability in weather patterns globally, and in temperate regions, there has been an increase in unseasonal conditions such as hail, snow, and night frosts that cause significant stress to plants and are potentially lethal. Low temperature freezing stress leads to significant and irreversible damage to cell membranes and oxidative stress, and causes physiological and metabolic changes that impact on plant growth and development [1]. The freezing injuries can be observed as necrosis in strawberry crown tissues. In some cases, the plants will recover from the injury, though with some loss in productivity. It has also been shown that a yield loss of up to 20% occurs before the damage is manifested as crown necrosis [2, 3]. In years with limited snow cover and low temperatures, entire fields can be destroyed. As such, cold stress and freezing tolerance in crop plants have become the focus of research efforts aiming to develop resilience to climate fluctuations, but it is also of significant importance for production in more northern regions of Europe, where extreme winter temperatures are encountered annually.

Strawberries (*Fragaria spp*.) are perennial plant species that are found growing naturally throughout the temperate regions of the world, including moderately high-altitude regions throughout their range [4]. Strawberries are cultivated globally, with production regions extending to the Nordic countries in Europe and Canada in North America. As such, strawberry species and those cultivated as crop plants, must survive extremely low temperatures during winter months. In major production areas in Norway, temperatures of –20 to –30 °C for several weeks are common. Without protection, either from a covering of snow or from frost protection covers, these temperatures can be devastating for the plants [5]. Low temperatures cause several physiological responses in strawberries including winter dormancy and autumn cold hardening, and in cold climates, species such as the diploid *F. vesca* have been shown to have adapted to low-temperature stresses. Genotypes from these regions are able to withstand much lower temperatures than accessions from more moderate climates [6, 7].

The diploid strawberry *F. vesca* is a model organism for studying development in perennial plant species, and a wealth of genetic resources are available to facilitate such studies [8-11]. In order to understand the genetic variation underlying winter survival in strawberry, a was conducted to characterise the low temperature stress tolerance of accessions of several diploid strawberry species [7]. In this study, accessions of *F. vesca* collected from regions with extreme climatic conditions, such as the north of Norway, were shown to have robust tolerance to freezing stress, whilst accessions from regions with milder climates, such as the subspecies *F. vesca* subsp. *californica* native to the west coast of the United States and *F. vesca* accessions naturalised in South American countries including Bolivia, were far more susceptible to freezing stress.

Here, we have investigated the genetic basis of freezing tolerance in *F. vesca*. A segregating mapping population derived from a cross between a freezing tolerant Norwegian *F. vesca* accession ‘Alta’ from the North of Norway, and a cold susceptible accession collected in South America was raised and phenotyped for freezing tolerance response under controlled and replicated laboratory conditions. A genetic map was produced using the genotyping by sequencing approach [12] and a significant quantitative trait locus associated with freezing tolerance was characterised. The physical interval underlying the QTL was interrogated, candidate genes were identified and their putative role in freezing tolerance was inferred from expression analyses. The possible role of these candidate genes in response to freezing stress is discussed.

## Materials and methods

### Plant material

Each genotype was propagated from runners to create a set of test plants of uniform size and developmental stage. The clonal plants were rooted and grown in 10 cm plastic pots containing a peat-based compost (90% peat, 10% clay), with the addition of 1:5 (v/v) of granulated perlite raised in a glasshouse for five weeks at 20 ± 2 °C and an 18-hour photoperiod. The plants were watered twice a week with a balanced nutrient solution containing 7.8 mmol N, 1 mmol P, and 4.6 mmol K per litre (used in 1:100 dilution).

### Freezing tolerance phenotyping

Prior to low temperature testing, the plants were acclimated at 2 °C and a 12-hour photoperiod for six weeks. Plants were watered with cold water as needed. After six weeks, plants were transferred to three programmable freezers where they were first kept at -2 °C for 12 hours. Subsequently the temperature was lowered by 2 °C/h until the target temperature was reached. The target temperature was held for 4 hours before raising it by 2 °C/h to 2 °C where they were maintained for a further 10 hours. Subsequently, the plants were kept overnight at room temperature, following which they were transferred to a greenhouse and maintained at 18 ± 2 °C with an 18-hour photoperiod for five weeks before survival (plants were observed to be dead or alive) was scored.

### Determination of suitable freezing temperatures for mapping population screen

The parental lines ‘Alta’ and ‘NCGR1363,’ selected for their differential response to freezing stress in a previous study [7] and an F_1_ hybrid line from the resultant cross ‘NCGR1363’ × ‘Alta’ were phenotyped to determine suitable temperatures at which to screen an F_2_ test population. The lines were subjected to freezing stress temperatures of -18 °C, -15 °C, -12 °C, -9 °C, -6 °C, and 0 °C following the procedure described above. For each temperature, 13 test plants of each genotype were screened.

### Freezing tolerance phenotyping of a segregating mapping population

The grandparental lines, F_1_ parent and an F_2_ mapping population consisting of 142 plants from the cross ‘NCGR1363’ × ‘Alta’ were propagated and subjected to cold tolerance screening. The optimal stress temperatures calculated from the ancestral germplasm screen (described above), to which the genotypes were subjected, were -5 °C, -8.5 °C, and -12 °C. Each F_2_ genotype was replicated nine times at each temperature, whilst the grandparental and F_1_ hybrid genotypes were replicated 18 times at each temperature. Five replicates of the entire experiment were performed at each temperature.

### Statistical data analysis

The analysis of the survival data (alive/dead) from both the experiment to determine optimal stress temperatures and the subsequent screening of the F_2_ mapping population, employed the following logistic model:

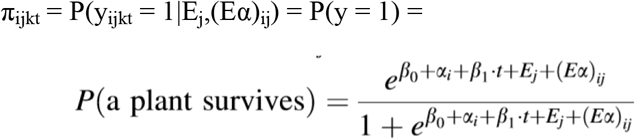

where π_ijkt_ is the observation [alive(1)/dead(0)] made on plant k from genotype *i*, in replicate *j*, exposed to temperature *t, β*_0_ is an unknown constant, *α*_i_ is the main effect of the genotype (*i*= 1, ….,n), *β*t is the coefficient that estimates the effect temperature (*t*) has on plant survival, *E*_j_ is the effect of replicate *j* (j= 1,…,5), (*Eα*)_ij_ is the interaction between the genotype *i* and replicate *j*.

The LT_50_ for genotype *i* was estimated as

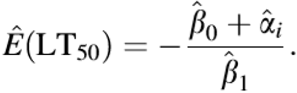

The *Glimmix* procedure in SAS® was used to implement this model for the F_2_-population while the similar calculations and the corresponding survival plot of the initial experiment (the parents and the hybrids) was drawn using R [13] following the code of Luke Miller (https://lukemiller.org/index.php/archive/).

### Mapping population development

The cross ‘NCGR1363’ (susceptible to low temperature stress) × ‘Alta’ (tolerant to low temperature stress) was performed at the NIBIO outstation in Kvithamar in Stjørdal, Norway. Hybrid nature of the plants were confirmed using a set of microsatellites [14]. A confirmed hybrid F_1_ plant from the cross was self-pollinated and a population of 142 segregating F_2_ seedlings (‘NCGR1363×Alta’) was raised for low-temperature stress tolerance phenotyping as detailed above and subsequent genetic map construction.

### Genotyping by sequencing (GBS) and SNP calling

DNA was extracted from emerging fresh leaf tissue of 12-week old plants of the ‘NCGR1363’ × ‘Alta’ mapping population, the F_1_ parental plant and the grandparental plants ‘NCGR1363’ and ‘Alta’ using the DNeasy Plant Minikit (Qiagen) and sample quality was determined using a QiAgility spectrophotometer (Qiagen). Samples were considered suitable for genotyping if they had a 260/280 ratio in the 1.8 to 2.0 range. DNA quantification was performed with a Qubit fluorometer against manufacturer-supplied standards (Thermo Scientific) and was normalised to 10 ng/ul. Genotyping data were generated from the grandparents, the F_1_ parent and the 142 progeny of the mapping population following the ‘Genotyping-by-Sequencing’ (GBS) protocol described by Elshire *et al*. [12]. Briefly, DNA was digested with the enzyme ApeKI and multiplexed fragment libraries were sequenced on an Illumina NextSeq 500 v2 sequencing machine, generating, on average, 1.5 million 75 bp single reads from each sample.

Demultiplexed raw reads from each sample were quality trimmed and aligned to the *F. vesca* v4.0 genome sequence using BWA-MEM version 0.7.12 [15] to create BAM files from which SNP variants were called using FreeBayes v1.0.2-16 [16] using the following specific parameters (--min-base-quality 10 --min-supporting-allele-qsum 10 --read-mismatch-limit 3 - -min-coverage 5 --no-indels --min-alternate-count 4 --exclude-unobserved-genotypes -- genotype-qualities --ploidy 2 or 3 --no-mnps --no-complex --mismatch-base-quality-threshold 10). Filtering of variants was performed with a GBS-specific rule set where read counts for a locus must exceed 8, minimum allele frequency across all samples must exceed 5% and genotypes must have been observed in at least 66% of samples.

### Linkage map construction and quantitative trait loci analysis (QTL) analysis

The resultant SNP data were used for mapping linkage map construction using JOINMAP 4.1 (Kyasma, NL). Following grouping, initial marker placement was determined using Maximum Likelihood with a minimum logarithm of odds (LOD) score threshold of 3.0, a recombination fraction threshold of 0.35, ripple value of 1.0, jump threshold of 3.0 and a triplet threshold of 5.0, and mapping distances were calculated using the Kosambi mapping function to produce individual linkage groups. Imputation was then performed following the protocol described by [17] and a second round of mapping using the parameters described above was performed to produce the final linkage map of the ‘NCGR1363’ × ‘Alta’ mapping progeny. The linkage map presented was plotted using MapChart 2.1.

QTL analysis was performed for the LT_50_ phenotypic data for the mapping progeny using interval mapping with MAPQTL 6.0 (Kyazma, NL) employing a step size of 1.0 cM, and the percentage phenotypic variance explained and associated LOD values were calculated. A LOD significance threshold of 3.2 was calculated following a permutation test with 10,000 reps, and was used to determine significance. The calculated LOD values were plotted with MapChart 2.3 [18] using the chart function.

### Functional variant identification and candidate gene analysis

Gene sequences for the predicted genes within the interval spanning the QTL identified on *Fvb2* of the *F. vesca* v4.0 genome sequence were extracted from the sequence data repository on the Genome Database for Rosaceae [19] and functionally annotated using OmicsBox (https://www.biobam.com/omicsbox/) running default parameters. Candidate genes within the QTL interval were identified based on the relevance to cold tolerance of their functional annotation.

### RNASeq analysis of QTL interval genes

Crown and leaf tissue from parental genotypes ‘Alta’ and ‘NCGR1363’ exposed to low temperature stress (2 °C for 42 d) and untreated plants (0 h controls) were ground to a powder under liquid nitrogen and RNA was extracted using the Spectrum Plant Total RNA Kit (Sigma) according to manufacturer’s instructions. Twenty-four libraries (two tissues types × two parental genotypes × two timepoints × three biological replicates) were prepared from RNA samples with a RIN (RNA integrity number) above 8 using the strand-specific TruSeq™ RNA-seq library (Illumina), and 150 bp paired-end read sequencing over three lanes of the Illumina HiSeq4000 sequencing platform was performed at the Norwegian Sequencing Centre, at the University of Oslo, Norway.

The fastq files generated were analyzed using FastQC (http://www.bioinformatics.babraham.ac.uk/projects/fastqc/) and TrimGalore (https://www.bioinformatics.babraham.ac.uk/projects/trim_galore/) was used for trimming of adapter sequences and low quality bases. Using the OmicsBox platform, trimmed reads were mapped to the predicted mRNA sequences from the *F. vesca* v.4.a2 reference genome annotation on the Genome Database for Rosaceae [19] using RSEM [20] and Bowtie2 [21].

Differentially expressed genes were identified using the edgeR (version 3.28.0; Robinson, McCarthy (22)) Bioconductor package that is integrated in the OmicsBox platform utilizing the following parameters: A false discovery rate (FDR) cut off of 0.05; generalized linear model (GLM) quasi-likelihood F-test; counts per million reads (CPM) cutoff of 0.5 in a minimum of 2 of 3 biological replicates; and sample normalization based on TMM (weighted trimmed mean of M-values) as recommended by the package user guide. Differentially expressed genes were then evaluated according to their functional annotation and those with a potential role in freezing tolerance as well as others observed to be highly differentially expressed were considered as potential candidates contributing to the identified QTL.

## Results

### Freezing tolerance phenotyping

The estimates of the temperatures at which 50% of the cohort of clones of a given genotype survived (LT_50_) along with their standard errors were -12.3 ± 0.25 °C for ‘Alta’ and -7.9 ± 0.28 °C for ‘NCGR1363’ which agreed with the results previously reported by Davik et al (2013). The F_1_ hybrid, ‘NCGR1363×Alta’ had an LT_50_ estimate of -10.7 ± 0.38 °C (Fig. 1) and estimated LT_50_ values were calculated for all 142 F_2_ progeny of the ‘NCGR1363’ × ‘Alta’ mapping population (see S1 Table). Data were plotted as frequency histograms (Fig. 2) and the LT_50_ estimates were used as phenotypes for quantitative trait locus detection.

**Fig 1.**
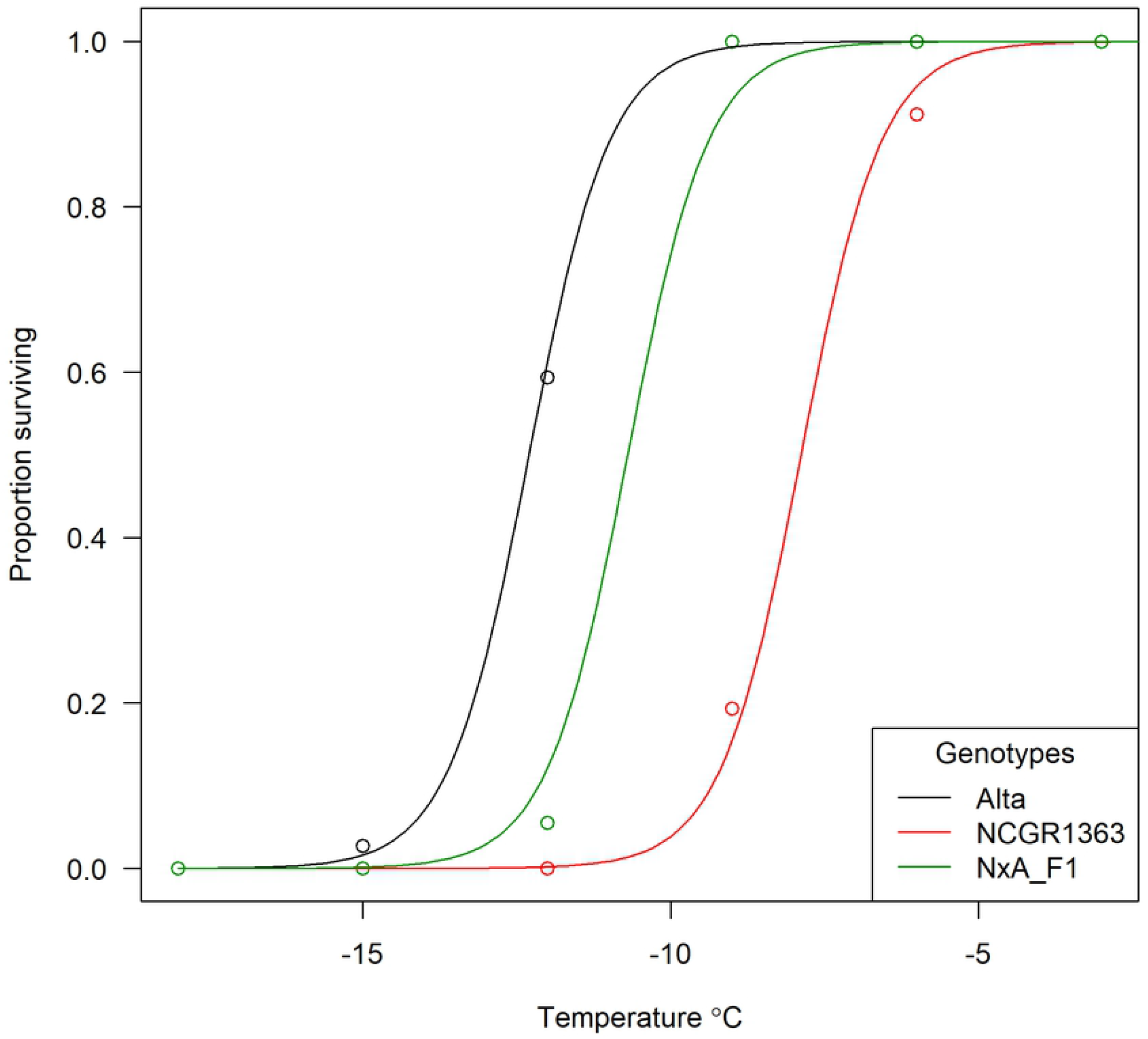
Alta, NCGR1363, and their hybrid show distinct freezing tolerance. Plot of the proportions of clones of *F. vesca* parental and F_1_ genotypes accessions surviving temperature stresses at -18 °C, -15 °C, -12 °C, -9 °C, -6, and 0 °C used to calculate the LT_50_ estimates (temperature at which 50% of the cohort of clones of a given genotype survived) for each accession.

**Fig 2.**
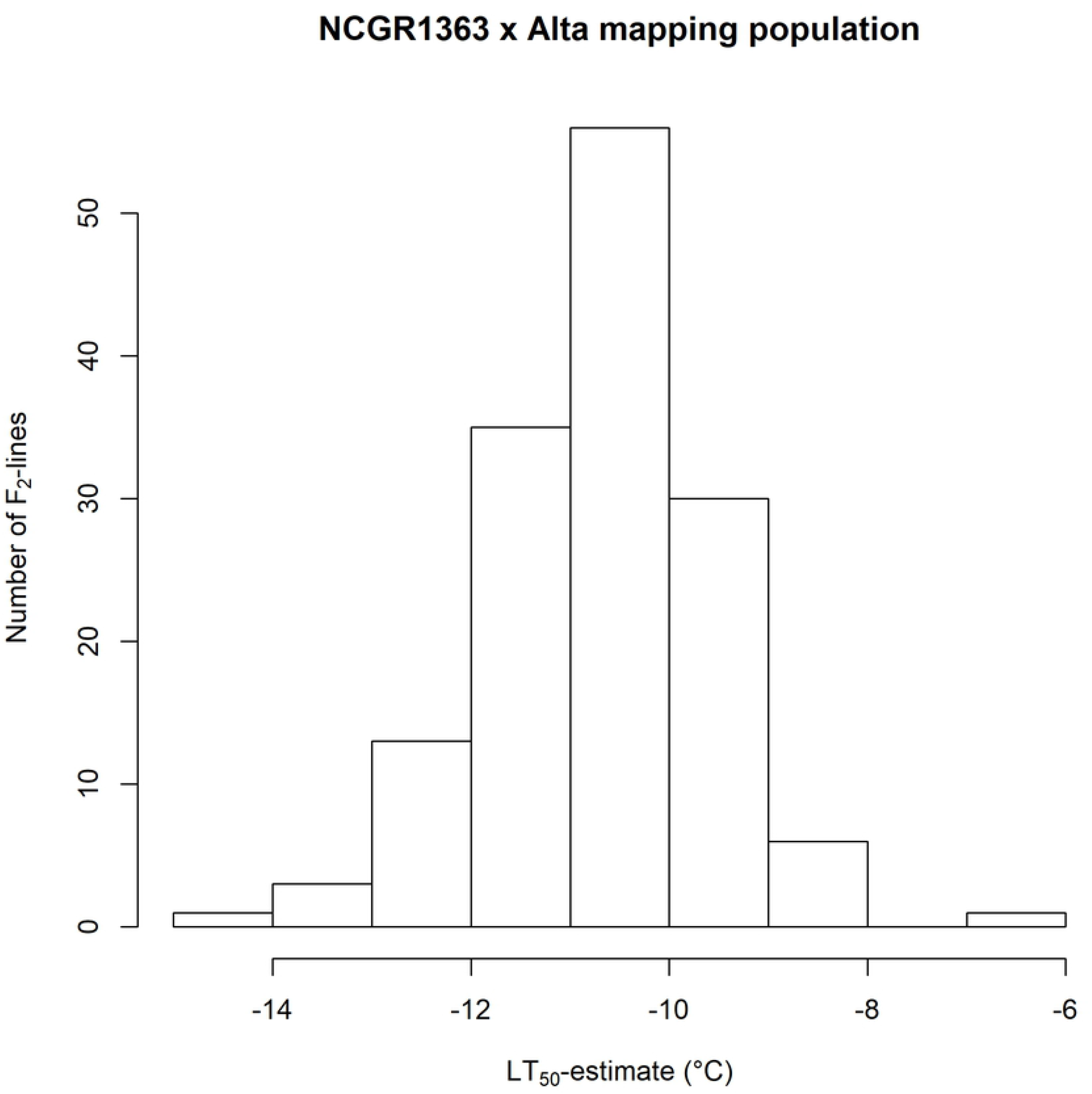
Progeny of ‘NCGR1363’ × ‘Alta’ show a “normal” distribution of freeze tolerance. Histogram of the LT_50_ estimated values calculated for the progeny of the ‘NCGR1363’ × ‘Alta’ mapping population (*n=*142). LT_50_ estimates are given in °C.

### Genotyping and linkage map construction

A total of 16,551 putatively polymorphic sequence variants were identified between the grandparental genotypes that were heterozygous in the F_1_ parent of the ‘NCGR1363’ × ‘Alta’ mapping progeny after data were analysed with the criteria described in the materials and methods. Of these, 3,294 clustered into one of seven discrete linkage groups corresponding to the seven pseudomolecules of the *F. vesca* v4.0 genome sequence following an initial round of linkage mapping. Following imputation and map construction, the seven resolved linkage groups contained a total of 2,918 markers at 851 loci and spanned a total genetic distance of 593.7 cM (Fig. 3; S2 Table), equating to a total physical distance on the *F. vesca* v4.0 genome sequence of 217.1 Mb. Linkage group 6 (*Fvb3*) was the longest, spanning 108.5 cM (36.7 Mb), whilst LG1 (*Fvb1*) was the shortest, spanning 57.6 cM (23.9 Mb).

**Fig 3.**
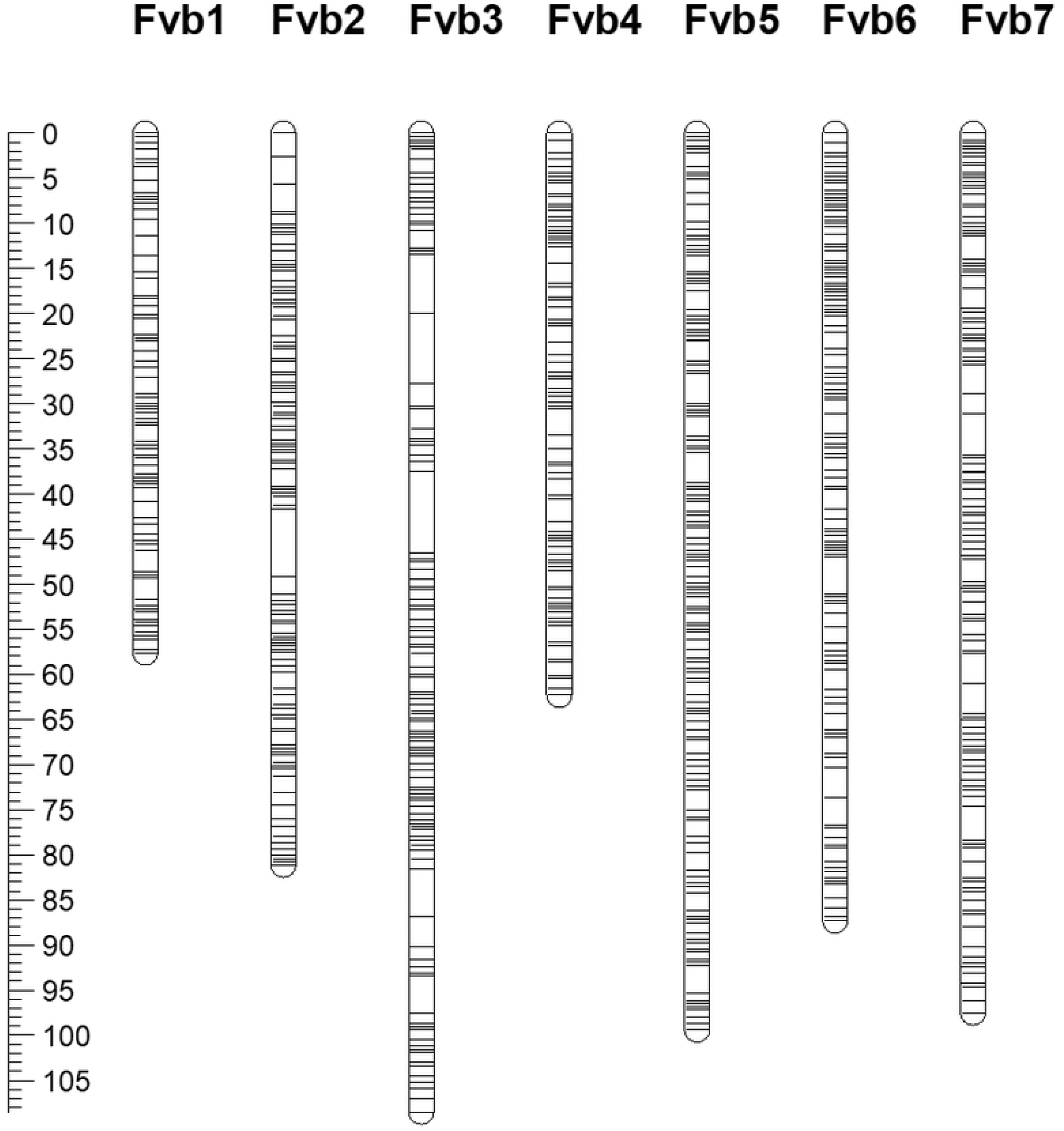
Genetic linkage map of the ‘NCGR1363’ × ‘Alta’ mapping population composed of seven linkage groups containing a total of 2,918 markers at 851 loci and covering a genetic distance of 593.7 cM. Linkage groups are named according to the seven *F. vesca* pseudochromosomes of the v4.0 genome sequence and genetic marker positions are given in cM.

### Quantitative trait loci analysis, functional variant identification, and candidate gene analysis

Following interval mapping implemented in MapQTL 4.0, a single significant QTL was identified on *Fvb*2 with a peak LOD score of 3.73 explaining 11.5% of the observed trait variance (Fig. 4). The most significant associations were with three SNP markers *Fvb2*_15730261 (10.4% observed variance explained LOD 3.36) and two co-segregating SNPs *Fvb2_*10601614 and *Fvb2*_10601635 (10.4% observed variance explained LOD 3.29) with physical positions at 10.6 Mb and 15.73 Mb on the *F. vesca* v4.0 genome sequence. As such, the QTL spanned an interval of 5,128,648 bp towards the proximal end of chromosome *Fvb2* of the *F. vesca* genome. The 5.1 Mb physical QTL interval on the *F. vesca* genome contained a total of 896 predicted genes, several of which have putative roles associated with tolerance to abiotic stresses including freezing. Among these were two gene predictions displaying high homology to *Alcohol Dehydrogenase 1* (*ADH1*; *FvH4_2g14760*.*1* and *FvH4_2g14750*.*1*), one encoding the dehydrin Early responsive to dehydration 10 (*ERD10*; *FvH4_2g16030*.*1*), two with homology to *PIP2* aquaporin genes (*FvH4_2g15440*.*1* and *FvH4_2g15450*.*1*), one with homology to *ascorbate oxidase* (*FvH4_2g16000*.*1*), one gene with homology to the glucose transporter *SWEET1* (*FvH4_2g14860*.*1*), an ABA-repressive AFP2-like regulator-encoding homolog (*FvH4_2g18440*.*1*), one encoding a gene with homology to a hAT dimerization domain-containing protein (abbreviated hereafter as hAT; *FvH4_2g12511*.*1*), a gene encoding a BYPASS1-like protein (B1L; *FvH4_2g13680*.*1*), one that encodes EXPANSIN-like A2 (EXLA2; *FvH4_2g16110*.*1*), a gene that encodes N-acetylserotonin O-methyltransferase (ASMT; *FvH4_2g15840*.*1*), a gene that encodes Ring and Domain of Unknown Function 2 (RDUF2; *FvH4_2g16170*.*1*), a serine/threonine protein-kinase CTR1-encoding gene (*FvH4_2g15800*.*1*), four gene predictions encoding NAC transcription factors (*FvH4_2g12690*.*1, FvH4_2g13330*.*1, FvH4_2g13320*.*1* and *FvH4_2g16180*.*1*), and a predicted gene (*FvH4_2g11510*.*1*) that encodes dynamin-related protein 3A (DRP3A) (S3 Table).

**Fig 4.**
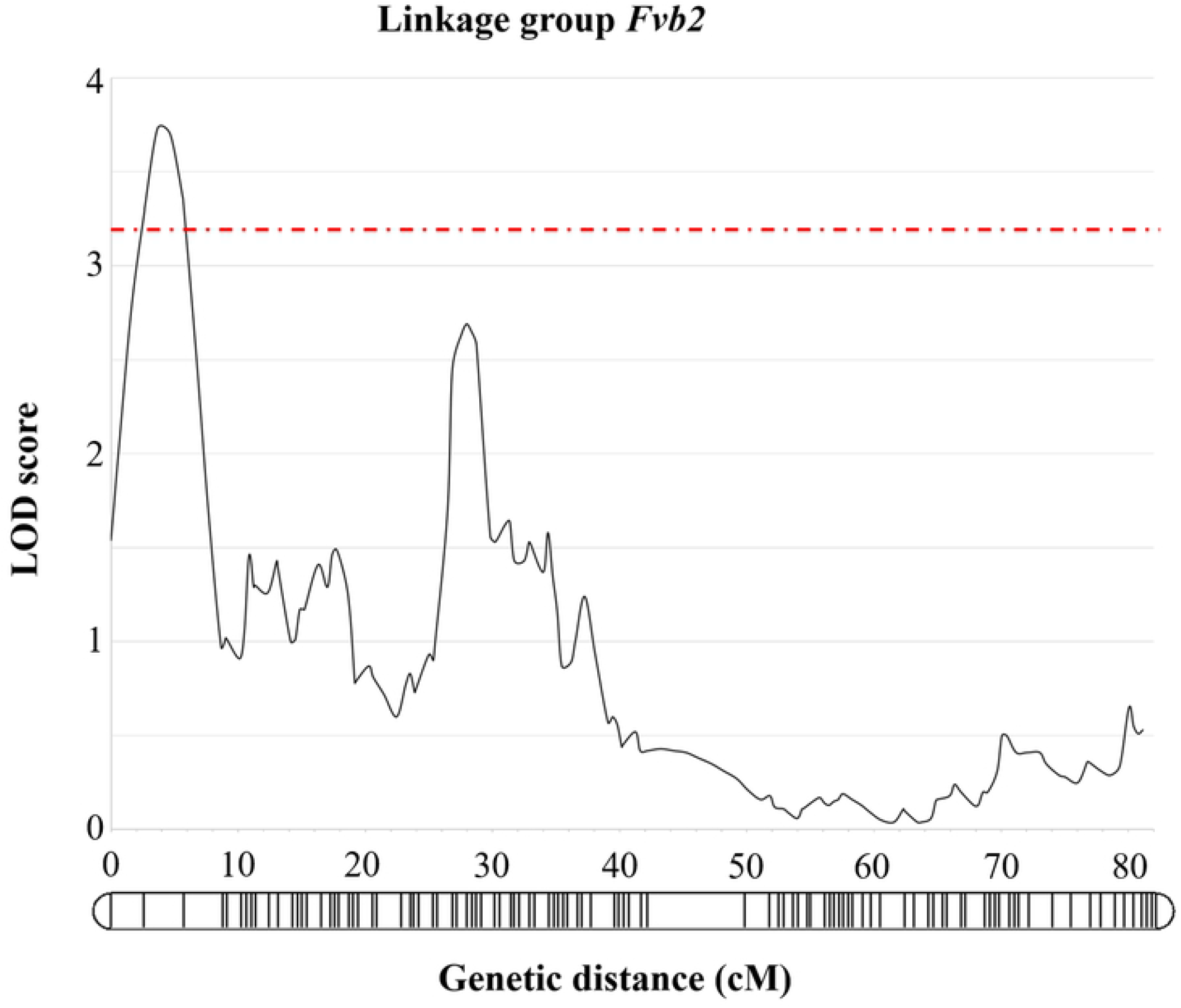
Significant QTL for LT_50_ identified on *Fvb2* of the ‘NCGR1363’ × ‘Alta’ mapping population. Genetic positions of the markers on *Fvb2* are shown along the linkage group and a LOD significance threshold of 3.2 is shown as a hashed line.

### Differential expression of candidate genes

Differential expression analysis of the 896 QTL-associated candidate genes in the leaves and crowns from ‘Alta’ and ‘NCGR1363’ parental genotypes treated for 0 H or 42 D at 2 °C revealed genotype-specific changes (fold change, FC) in transcript accumulation in response to low temperature treatment as well as expression differences (fold difference, FD) between genotypes prior to- (i.e. basal transcript levels), and after treatment (S4 Table). Fourteen of the candidate genes identified in the QTL interval that were putatively associated with plant response to temperature and/or osmotic stresses, exhibited parental genotype differences in transcript accumulation either prior to treatment or in response to cold treatment, sometimes both (S4 Table). Candidate genes showing genotype-specific cold-responsive expression differences primarily in crown tissue were DRP3A (*FvH4_2g11510*), SWEET1-like (*FvH4_2g14860*.*1*), hAT (*FvH4_2g12511*.*1*), BYPASS1-like (*FvH4_2g13680*.*1*), ASMT (*FvH4_2g15840*.*1*) and NAC029 (*FvH4_2g16180*.*1*). The hAT, BYPASS1-like, ASMT, and NAC029 homologues exhibited a greater cold-responsive down regulation in ‘Alta’ crowns when compared to ‘NCGR1363,’ while SWEET1-like showed a greater cold-induced increase in ‘Alta’ when compared to ‘NCGR1363’, and the t3 isoform of *FvH4_2g11510*.*1* (*DRP3A*) displayed a FD of greater than -1600 in cold-treated crowns of ‘NCGR1363’ when compared to ‘Alta.’ Additional differentially-expressed genes were identified with a role in abiotic plant stress, including those encoding a vacuolar iron transporter (*FvH4_2g13960*.*1*), an acyl-[acyl-carrier-protein] (*FvH4_2g14690*.*1*), two MAD3/HGMR homologs (*FvH4_2g17051*.*1* and *FvH4_2g17060*.*1*), a major facilitator superfamily protein-encoding gene (*FvH4_2g15730*.*1*) and flavin-containing monooxygenase 1 (*FvH4_2g16430*.*1*) (Supplementary table S3, section B). With the exception of DRP3A, the directions of LTS-induced FC effects appear to nearly offset disparate basal expression levels between genotypes.

Candidate genes displaying genotype-specific cold-responsive expression differences primarily in leaf tissue, and basal pre-LT-treatment levels in crowns were a PIP2;7 homolog isoform (*FvH4_2g15440*.*t3*), CTR1 (*FvH4_2g15800*.*1*), a gene encoding a homolog of EXPLA2 (*FvH4_16110*.*1*) and RDUF2 (*FvH4_2g16170*.*1*) that also exhibited genotype-specific differences in LT-treated crowns, with much lower expression in ‘NCGR1363’ compared to ‘Alta.’ Additionally, further candidate genes without a clear association to low temperature stress were identified, including two laccases (*FvH4_2g12570*.*1* and *FvH4_2g12620*.*1*), an ABR1-like transcription factor (*FvH4_2g13240*.*1*) carbonic anhydrase 2-like homolog (*FvH4_2g14190*.*1*), OXI1 kinase (*FvH4_2g18040*.*1*) and a lysozyme D-like homolog (*FvH4_2g18210*.*1*).

A third group contained two genes differentially expressed between genotypes primarily in leaves, both in pre-LT-treated and LT-treated plants. These were a homolog of a known cold response pathway gene ninja-family AFP2-like (*FvH4_2g18440*.*1*) and a P450 87A3-like homolog (*FvH4_2g16500*.*1*), not previously known for a role in temperature stress. These genes showed opposite regulation patterns with greater AFP2-like transcript accumulation in cold-treated ‘Alta’ leaves, while ‘NCGR1363’ exhibited vastly higher transcript levels, primarily in leaves of non-LT- and LT-treated plants.

## Discussion

One of the aims of a previous investigation was to identify parental genotypes that would be suitable for studying the genetics of freezing tolerance in *F. vesca* [7]. Here, the genotypes displaying the greatest differences in freezing tolerance were crossed and from the selfing of the resultant F_1_ progeny, a mapping population segregating for freezing tolerance was raised and phenotyped. A freezing tolerance QTL identified on chromosome *Fvb2* of the *F. vesca* genome spanned a physical interval of 5.1 Mb and contained 896 gene predictions encoding proteins including ADH1, ERD10, PIP2 aquaporins, ascorbate oxidase, a hAT dimerization domain-containing protein, CTR1 kinase, B1L, ASMT, EXPLA2, transcription factor RDUF2, a ninja-family AFP2-like transcription factor and two NAC transcription factors, all of which have been reported to have putative roles in plant temperature or osmotic stress response and, for some, freezing tolerance. Here we discuss the candidate genes identified in the context of freezing tolerance in the ‘NCGR1363’ × ‘Alta’ *F. vesca* mapping population.

### Alcohol dehydrogenase

An increase in the production of C1 to C9 alcohols in plants enhances membrane fluidity which prevents phase transition occurring in plant cell membranes, thus promoting greater tolerance to freezing stress in plants [23]. Alcohol dehydrogenase (ADH) genes play a role in the production of C1 to C9 alcohols and have been shown to be cold-induced in *Arabidopsis* and cereal crops [24]. More recently, Song [25] reported that in *Arabidopsis*, ADH1 was significantly upregulated in response to cold treatment and the ADH1 knockout mutants they screened showed lower basal freezing tolerance than wild-type plants, and a higher percentage of ion leakage after freezing treatment, suggesting a pivotal role for ADH1 in the protection of plasma membranes and thus in freezing stress tolerance. An ADH1 homologue was the first protein-encoding gene to be completely sequenced in cultivated strawberry [26], was the first gene sequence to be genetically mapped in *Fragaria* [27], and has been used to infer phylogenetic relationships in the genus [28]. Koehler *et al*. [29] reported a strong correlation between ADH levels and cold tolerance in the cultivated strawberry *F. ×ananassa* and similarly, Davik *et al*. [7] demonstrated that LT_50_ was strongly correlated (r = -0.86) with ADH protein levels. Davik *et al*. [7] also reported that ADH levels were very low in *F. vesca* control crowns, but strongly induced in cold-treated crowns, with up to a 200-fold increase in ADH protein levels observed after 42 days of cold treatment in accessions that were shown to be highly tolerant to freezing stress. The authors concluded that ADH likely contributes to cold hardiness in *F. vesca*.

The ADH1 homologue first mapped by Davis and Yu [27] is located at 12,948,939 bp on chromosome *Fvb2* of the *F. vesca* genome, placing it within the mapping interval of the QTL identified in this investigation for freezing tolerance in *F. vesca* ‘Alta.’ Both ADH homologs were expressed at higher basal levels in ‘NCGR1363’ crowns but appeared to show a lower cold-induced increase, especially for *FvH4_2g14760*.*1*, in leaves in comparison with ‘Alta’. Neither of these homologs displayed the cold-induced increases at the transcript level in crowns that was previously observed for immunoreactive ADH proteins [7]. The higher expression of ADH transcripts in leaves of ‘Alta’ correlates well with its lower LT_50_.

### Dehydrins (FvH4_2g16030.1, FH4_2g09610.1)

The expression of dehydrins has previously been reported to be highly correlated to cold-stress tolerance in cultivated strawberry [30]. More recently, Koehler et al [29] performed gene expression and proteomic profiling of the commercial cultivars ‘Jonsok’ and ‘Frida’ following cold exposure and demonstrated that the transcript levels of two dehydrin-like genes, a COR47-like and a XERO2-like gene were strongly correlated with cold stress. The authors speculated that the strong increase in observed levels of a dehydrin protein identified through a one-dimensional electrophoresis western-blot that used an anti-K peptide diagnostic for dehydrin was the XERO2-like dehydrin. In a proteomics study of *F. vesca*, dehydrin accumulation was observed following 14 days of cold treatment, with higher levels of seven distinct dehydrins accumulating after 42 days cold treatment [7]. Further, an examination of the natural variation in cold/freezing tolerance in diploid *Fragaria* genotypes showed a strong correlation of plant survival with the expression of total dehydrins [7]. However, due to non-specificity of *Arabidopsis* dehydrin antibodies, the authors were unable to determine which specific dehydrins accumulated in the study.

The physical region spanning the QTL identified in this investigation contains a predicted dehydrin gene (*FvH4_2g16030.1)* with strong homology to the acidic class of dehydrins (exemplified by the Arabidopsis ERD10, ERD14, and COR47). The ERD (early responsive to dehydration stress) genes, while first identified as rapidly upregulated in response to dehydration in Arabidopsis, were subsequently observed to be upregulated by cold and other stresses [31]. ERD10, an ABA-dependent dehydrin, was characterised in *Brassica napus* [32] where it was shown to be induced in leaf tissue by cold stress, and subsequently it was also reported to be induced in response to cold stress in *Arabidopsis* [33, 34]. Strong evidence in strawberry [30] and other plants demonstrate that over- or trans-expression of dehydrins increase cold and other stress tolerances [35-38]. However, the ERD-encoding homolog *FvH4_2g16030.1* did not show expression in the material utilized for RNASeq analysis of QTL candidate genes.

### Plant intrinsic proteins, aquaporin (FvH4_2g15440.1, FvH4_2g15450.1)

Aquaporins are a highly conserved group of membrane proteins which help transport water across biological membranes and are known as major intrinsic proteins. The plasma-membrane intrinsic proteins (PIPs) are a class of aquaporins that are highly responsive to environmental stimuli and have roles in various physiological functions including response to drought stress [39]. The PIP gene family, comprising 13 genes in *Arabidopsis thaliana* have been shown to be expressed under various abiotic conditions including drought, cold, and high salinity stress, as well as abscisic acid (ABA) treatment [40]. In the study of [40], PIP2;5 was shown to be up-regulated by cold stress, while most of the other members of the family were down-regulated. Similarly, in a proteomics study of cold stress in banana species (*Musa* spp. ‘Dajiao’ and ‘Cavendish’), the abundance of aquaporins significantly increased after 3 hours of cold stress in ‘Dajiao’ seedlings [41] and the authors concluded that the aquaporins MaPIP1;1, MaPIP1;2, MaPIP2;4, MaPIP2;6, and MaTIP1;3 were all involved in decreasing lipid peroxidation and maintaining leaf cell water potential in cold stressed seedlings, which were likely the cellular adaptations responsible for increased cold tolerance of ‘Dajiao’ over ‘Cavendish’ seedlings. A total of ten PIP aquaporins have previously been reported in the genome of *F. vesca* [39], where diurnal expression was observed in the transcript levels of three of the characterised genes. More recently, substrate-specific expression profiles were shown for aquaporins in the cultivated strawberry *F. ×ananassa* [42], suggesting functional specialisation of aquaporins within the same class. Two predicted genes identified within the QTL interval displayed high homology to PIP2 aquaporins and could thus play a role in freezing-stress tolerance in *F. vesca*. The RNASeq analyses performed here revealed the t3 mRNA isoform of *FvH4_2g15440* was expressed at higher levels in pre-LT-treated crowns and displayed a greater LT stress induction in leaves of ‘Alta’ when compared with ‘NCGR1363.’ As with the ADH homologs in the QTL interval, involvement of this single PIP2;7 isoform, one of five from this gene, in contributing to freezing tolerance would be predicated with such a role being exercised primarily in leaves.

### NAC transcription factors (FvH4_2g16180.1, FvH4_2g12690.1, FvH4_2g13330.1, FvH4_2g13320.1)

NAC transcription factors are one of the largest families of transcription factors in plants and have been implicated in enhancing tolerance to various abiotic stresses including drought, high salinity and cold, in a number of plants [43, 44]. In apple (*Malus pumila*) a close relative of *Fragaria* in the Rosaceae family, a NAC transcription factor MdNAC029 was shown to be a negative modulator of the cold stress response, directly repressing the expression of two C-repeat binding factors, MdCBF1 and MdCBF4, which are regarded key regulators of the plant response to cold stress [45]. Similarly, the role of NAC transcription factors in cold-stress response was studied in *Prunus mume* another member of the Rosaceae family, and 113 PmNAC genes were identified and characterised [46]. Seventeen of the genes identified were highly up-regulated in stem tissue during cold temperature stress during winter. Further analysis of a subset of 15 NAC genes showed that they were up and down-regulated in response to low-temperature treatment and were suggested to be putative candidates for regulating freezing resistance in the species. Within the freezing tolerance QTL identified in this investigation, candidate genes were identified with homology to three NAC transcription factors, NAC017, shown to negatively regulate drought-stress responses in *Arabidopsis* [47], NAC082, reported to be a ribosomal stress response mediator [48] and a homologue of NAC029, involved in cold-stress in apple [45] and upregulated in response to cold stress in *Gossypium barbadense* [49]. In this investigation, NAC029 showed a greater down-regulation in ‘Alta’ crowns when compared to ‘NCGR1363,’ however the latter showed a much lower pre-LT basal crown expression of this gene, and no significant expression differences between parents were detectable in cold-treated crowns. Despite this, the downregulation of NAC029 in response to cold is consistent with a role of this protein as a negative regulator of cold stress response in *Fragaria* as was observed in apple [45].

### Ascorbate oxidase (FvH4_2g16000.1)

Abiotic stress induces excess reactive oxygen species (ROS) which cause oxidative stress in plants resulting in damage to lipids, DNA, RNA and proteins. ROS detoxification systems are needed to protect plant cells against the toxic effect of these species [50, 51]. The ascorbate/glutathione pathway can ameliorate the oxidative stress. Ascorbate redox status in the cell wall is regulated by the apoplastic ascorbate oxidase (AO) where it catalyses oxidation of ascorbate to monodehydroascorbate (MDHA). The short-lived MDHA may then be reduced by a membrane-associated cytochrome B or disproportionate to ascorbate and dehydroascorbate (DHA). The increased transport of DHA into the cell would be expected to lead to an alteration of the overall redox status of ascorbate, decreasing its ability to provide antioxidative support. This possibility is consistent with the observation that AO-deficient RNAi antisense mutants are more tolerant to salt and oxidative stresses than WT while overexpressing plants are susceptible to these treatments [52-54]. However, the correlation between AO expression levels in the parental genotypes, at least concerning crowns, and their LT_50_ phenotypes contrasts with the previously observed effects relating to tolerance to salt and osmotic stresses.

### Other QTL-related candidate genes showing interparental differential expression

In addition to candidate genes with an identifiable role in freezing tolerance from previous studies, several genes within the QTL interval, whilst lacking clear association with low temperature stress tolerance in the Rosaceae, have been previously connected with temperature and/or osmotic stress response in other plants species.

### RDUF2 homolog (FvH4_2g16170.1)

RDUF2 is an E3 ubiquitin ligase whose expression in *Arabidopsis* was shown to be enhanced by salt, drought and ABA-treatment, and a knock-out mutant of this AtRDUF2 exhibited markedly reduced tolerance to drought stress [55]. RDUF2 is likely part of ABA-mediated positive regulation of drought responses in plants. In both pre- and post-LT treated crowns and leaves, ‘NCGR1363’ expressed the RDUF2 homolog (*FvH4_2g16170.1*) at levels 200- to nearly 900-fold lower than in ‘Alta’, making it a strong candidate gene in the identified QTL interval.

### CONSTITUTIVE TRIPLE RESPONSE1 (CTR1; FvH4_2g15800.1)

The CONSTITUTIVE TRIPLE RESPONSE1 (CTR1), a Raf-like Ser/Thr protein kinase, is a negative regulator that inhibits ethylene signal transduction [56, 57] which functions as an essential upstream positive regulator of EIN3 in ethylene signalling [58]. Shi *et al*. [59] demonstrated that both a *ctr1* mutant and an *EIN3*-over-expressing line displayed enhanced freezing tolerance in Arabidopsis. While the *CTR1* homologue *FvH4_2g15800.1* showed no cold-induced expression changes in parental crown tissue, the basal level in ‘NCGR1363’ was over 5-fold lower than ‘Alta.’ The data presented here showed a cold-induced fold change increase in CTR1 transcript accumulation in ‘Alta’ leaves while changes in ‘NCGR1363’ leaves were not significant.

### EXPANSIN-like A2 (EXLA2; FvH4_2g16110.1)

The Arabidopsis EXPANSIN-like A2 (*EXLA2*) gene was first characterised by its regulation and role in responses to biotic stress, namely infections with the necrotrophic pathogen *Botrytis cinerea, Pseudomonas syringae* pv. tomato, and the necrotrophic fungus *Alternaria brassicicola* [60]. Expansins cause loosening and extension of the cell wall, possibly by disruption of noncovalent bonding between cellulose microfibrils and matrix glucans [61]. The *exla2* mutant described by Abuqamar *et al*. [60] exhibited hypersensitivity to salt and cold stress. The *exla2* homologue in this study (*FvH4_2g16110*.*1*) displayed lower cold-responsive changes in transcript accumulation in both crowns and leaves of ‘NCGR1363’ compared with ‘Alta.’

### BYPASS1-like (FvH4_2g13680.1)

BYPASS1-like is a DUF793 family protein rapidly induced under cold treatment in *Arabidopsis* and is thought to enhance freezing tolerance in plants through stabilizing CBF3 and ensuring normal *CBF* and CBF target gene expression [62]. While ‘NCGR1363’ exhibited a 6.4-fold lower basal expression of a B1L homolog (*FvH4_2g13680.1*) in crowns compared to ‘Alta,’ transcript levels did not change in response to 42 d cold treatment, whereas ‘Alta’ reduced B1L transcript levels 7.4-fold to levels closely matching ‘NCGR1363’ expression following cold treatment.

### SWEET1-like homologue (FvH4_2g14860.1)

Expression of the SWEET1-like homologue *FvH4_2g14860.1* was 9-fold lower in crown tissue of cold-treated ‘NCGR1363’ than in ‘Alta.’ SWEET proteins are a family of oligomeric sugar transporters in plants, and in *Arabidopsis*, disruption of AtSWEET11 and AtSWEET12, normally down-regulated in response to cold stress, display increased freezing tolerance in an *AtSWEET11 AtSWEET12* double mutant [63]. It is conceivable that the higher cold-responsive expression of *SWEET1-like* in ‘Alta’ crowns therefore could sequester the *Fragaria* homolog of SWEET11 in a complex to further lower the functional levels of this protein during cold stress.

### hAT dimerization domain-containing protein (FvH4_2g12511.1)

The hAT transposon superfamily encodes transposase proteins harbouring dimerisation domains [64]. The gene *FvH4_2g12511.1* in *F. vesca* encodes a homolog hAT dimerization domain-containing protein. Transcript accumulation from this gene was over 300-fold lower in cold-treated crown tissue in ‘Alta’ compared with the less cold-tolerant ‘NCGR1363.’ Although it is not known whether this gene harbours functions unrelated to transposon activity, it has been shown that conserved genes derived from transposable elements are associated with abiotic stress phenotypes in *Arabidopsis* [65]. If it functions to increase transposition, elevated expression in ‘NCGR1363’ in response to cold stress would likely be detrimental.

### AFP2-like homolog (FvH4_2g18440.1)

AFP2 is one of four members of a family of ABI FIVE binding proteins in *Arabidopsis*. Knock-down *afp2-1* mutant plants were shown to be hypersensitive to salt, glucose and osmotic stress, but only mildly hypersensitive to ABA [66]. In addition to induction of stomatal closure and tolerance of drought, and salt stress, vegetative responses to ABA include cold stress tolerance (reviewed in Leung and Giraudat (67)). Significantly lower expression of the ninja-family AFP2-like homolog (*FvH4_2g18440.1*) in ‘NCGR1363’ leaves compared with ‘Alta’ suggests a possible role in the freezing tolerance observed here.

### ASMT homolog (FvH4_2g15840.1)

Phyto-melatonin, synthesized by ASMT, is postulated to mediate plant stress responses by counteracting stress-induced ROS [68]. Direct evidence of the cold-tolerance promoting properties of melatonin, also in the Rosaceae, stem from effects of exogenous application (e.g., Gao, Lu (69)). The ASMT homolog *FvH4_2g15840.1* contained in the QTL interval displayed a 2.5 lower basal expression in ‘NCGR1363’ crowns compared to ‘Alta’, and both exhibited a downregulation of ASMT transcripts in cold-treated crowns, the extent of which was over 100 times greater in ‘Alta’. LT-induced ASMT downregulation appears inconsistent for melatonin playing a role in enhanced cold tolerance in ‘Alta’, but may indicate that higher basal, pre-LT expression is the mechanism through which melatonin prepares plants for improved tolerance to low temperatures; certainly, the effects of melatonin on cold tolerance in the Rosaceae [69] and Arabidopsis [70] are based solely on pre-chilling treatments.

## Conclusions

Freezing tolerance is a quantitative complex trait with numerous genetic factors and a strong environmental component contributing to its expression. In this investigation, we identified a significant QTL that explained 10.4% of the phenotypic variance observed in a *F. vesca* mapping population which was located in a wide physical interval on *Fvb2* of the *F. vesca* v4.0 genome sequence. The physical interval was relatively large, spanning 5.1 Mb, and gene expression studies of the crowns and leaves of parental cultivars during cold-stress highlighted several potential candidate genes within the interval that could be responsible for the variation observed in freezing tolerance of the ‘NCGR1363’ × ‘Alta’ progeny.

Significant interparental differential expression levels of several of the genes reported here, along with previous evidence for roles for many of them in cold- and freezing-temperature stress responses, identified them as good candidates for the control of the effects of freezing tolerance at the QTL identified in this investigation. In order to determine the causal genetic factor for the freezing tolerance observed, further functional annotation and characterisation of the candidate genes identified will need to be performed, including the identification of causal genetic variants in the grand-parental, parental and progeny lines of the ‘NCGR1363’ × ‘Alta’ mapping population and additional studies of candidate genes expression at further time-points during challenge with low-temperature stress which were beyond the scope of this current investigation. A greater knowledge of the genetic elements influencing tolerance to low-temperature stress and freezing could help develop new strawberry varieties adapted to growing environments at higher latitudes and capable of surviving in extreme winter conditions in years with no snow cover.

## Acknowledgements

The research has been supported by grants to JD, RCW, SKR, and MKA from the Norwegian Resarch Council (#179466/I11 and #244658/E50)

## Author contributions

JD conceived the study, performed the experimentation, analysed data and interpreted the results, and authored the manuscript; RCW, RGN, and PEG performed experimentation, analysed data and authored the manuscript; SKR and MKA authored the manuscript; DJS analysed data, interpreted results and authored the manuscript. All authors approved the manuscript.

## References

1. Kazemi-Shahandashti S-S, Maali-Amiri R. Global insights of protein responses to cold stress in plants: Signaling, defence, and degradation. J Plant Physiol. 2018;226:123–35. doi: https://doi.org/10.1016/j.jplph.2018.03.022.

2. Daugaard H. Winter hardiness and plant vigor of 24 starwberry cultivars grown in Denmark. Fruit Varieties Journal. 1998;52(3):154–7.

3. Nestby R, Bjørgum R, Nes A, Wikdahl T, Hageberg B. Winter cover affecting freezing injury in strawberries in a coastal and continental climate. J Hort Sci Biotech. 2000;75(1):119–25.

4. Staudt G. Systematics and geographic distribution of the American stawberry species. Taxonomic studies in the genus Fragaria (Rosaceae:Potentilleae): Universtity of California; 1999 1999.

5. Davik J, Daugaard H, Svensson B. Strawberry Production in the Nordic countries. Adv Strawb Prod. 2000;19:13–8.

6. Sønsteby A, Heide OM. Environmental Regulation of Dormancy and Frost Hardiness in Norwegian Populations of Wood Strawberry (*Fragaria vesca* L.). In: Nestby, R (ed) Plant Science and Biotechnology in Norway Europ J Plant Sci Biotech 2011. p. 42–8.

7. Davik J, Koehler G, From B, Torp T, Rohloff J, Eidem P, et al. Dehydrin, alcohol dehydrogenase, and central metabolite levels are associated with cold tolerance in diploid strawberry (*Fragaria* ssp.). Planta. 2013;237:265–77.

8. Shulaev V, Sargent DJ, Crowhurst RN, Mockler TC, Folkerts O, Delcher AL, et al. The genome of woodland strawberry (*Fragaria vesca*). Nat Genet 2011. p. 109–16.

9. Edger PP, VanBuren R, Colle M, Poorten TJ, Man Wai C, Niederhuth CE, et al. Single-molecule sequencing and optical mapping yields an improved genome of woodland strawberry (Fragaria vesca) with chromosome-scale contiguity. GigaScience. 2018;7:1–7.

10. Hawkins C, Caruana J, Li J, Zawora C, Darwish O, Wu J, et al. An eFP browser for visualizing strawberry fruit and flower transcriptomes. Hort Res. 2017;4:17029.

11. Kang C, Darwish O, Geretz A, Shahan R, Alkharouf N, Liu Z. Genome-Scale Transcriptomic Insights into Early-Stage Fruit Development in Woodland Strawberry *Fragaria vesca*. The Plant Cell. 2013;25(6):1960–78. doi: 10.1105/tpc.113.111732.

12. Elshire RJ, Glaubitx JC, Sun Q, Poland JA, Kawamoto K, Buckler ES, et al. A robust, simple genotyping-by-sequencing (GBS) approach for high diversity species. PLoS ONE. 2011;6:e19379.

13. R Core Team. R: A language and programming environment for statistical computing. 2016.

14. Govan CL, Simpson DW, Johnson AW, Tobutt KR, Sargent DJ. A reliable multiplexed microsatellite set for genotyping *Fragaria* and its use in a survey of 60 *F. x ananassa* cultivars. Mol Breed. 2008;22(4):649–61.

15. Li H. Aligning sequence reads, clone sequences and assembly contigs with BWA-MEM. arXiv:13033997v1 [q-bioGN]. 2013.

16. Garrison E, Marth G. Haplotype-based variant detection from short-read sequencing. arXiv. 2012;arXiv:1207.3907[q-bio.GN].

17. Ward JA, Bhangoo J, Fernandez-Fernandez F, Moore P, Swanson JD, Viola R, et al. Saturated linkage map construction in *Rubus idaeus* using genotyping by sequencing and genome-independent imputation. BMC Genomics. 2013;14.

18. Voorrips RE. MapChart: Software for the Graphical Presentation of Linkage Maps and QTLs. J Hered. 2002;93(1):77–8. doi: 10.1093/jhered/93.1.77.

19. Jung S, Lee T, Cheng C-H, Buble K, Zheng P, Yu J, et al. 15 years of GDR: New data and functionality in the Genome Database for Rosaceae. Nucleic Acids Res. 2018;47:D1137–D45.

20. Li B, Dewey CN. RSEM: accurate transcript quantification from RNA-Seq data with or without a reference genome. BMC Bioinformatics. 2011;12(1):323. doi: 10.1186/1471-2105-12-323.

21. Langmead B, Salzberg SL. Fast gapped-read alignment with Bowtie 2. Nat Methods. 2012;9(4):357–9. doi: 10.1038/nmeth.1923.

22. Robinson MD, McCarthy DJ, Smyth GK. edgeR: a Bioconductor package for differential expression analysis of digital gene expression data. Bioinformatics. 2010;26(1):139–40. Epub 2009/11/17. doi: 10.1093/bioinformatics/btp616. PubMed PMID: 19910308; PubMed Central PMCID: PMCPMC2796818.

23. Thomashow MF. PLANT COLD ACCLIMATION: Freezing Tolerance Genes and Regulatory Mechanisms. Annu Rev Plant Physiol Plant Mol Biol. 1999;50(1):571–99. doi: 10.1146/annurev.arplant.50.1.571. PubMed PMID: 15012220.

24. Lindlöf A, Bräutigam M, Chawade A, Olsson B, Olsson O. Identification of Cold-Induced Genes in Cereal Crops and *Arabidopsis* Through Comparative Analysis of Multiple EST Sets. In: Hochreiter, S and Wagner, R (eds) Bioinformatics Research and Development - First International Conference BIRD ‘07, LNBI Vol 4414, Springer, Berlin/Heidelberg2007. p. 48–65.

25. Song Y, Liu L, Wei Y, Li G, Yue X, An L. Metabolite Profiling of *adh1* Mutant Response to Cold Stress in *Arabidopsis*. Frontiers in Plant Science. 2017;7(2072). doi: 10.3389/fpls.2016.02072.

26. Wolyn DJ, Jelenkovic G. Nucleotide sequence of an alcohol dehydrogenase gene in octoploid strawberry (*Fragaria* x *ananassa* Duch). Plant Mol Biol. 1990;14(5):855–7.

27. Davis TM, Yu H. A linkage map of the diploid strawberry, *Fragaria vesca*. J Hered. 1997;88:215–21.

28. DiMeglio LM, Staudt G, Yu H, Davis TM. A phylogenetic analysis of the genus *Fragaria* (Strawberry) using intron-containing sequence from the ADH-1 gene. PLoS ONE2014. p. e102237.

29. Koehler G, Wilson RC, Goodpaster JV, Sønsteby A, Lai X, Witzmann FA, et al. Proteomic study of low temperature responses in strawberry cultivars (*Fragaria* x *ananassa* Duschesne) that differ in cold tolerance. Plant Physiol 2012. p. 1787–805.

30. Houde M, Dallaire S, N’Dong D, Sarhan F. Overexpression of the acidic dehydrin WCOR410 improves freezing tolerance in transgenic strawberry leaves. Plant Biotechnol J 2004. p. 381–7.

31. Kiyosue T, Yamaguchi-Shinozaki K, Shinozaki K. Cloning of cDNAs for genes that are early-responsive to dehydration stress (ERDs) in*Arabidopsis thaliana* L.: identification of three ERDs as HSP cognate genes. Plant Mol Biol. 1994;25(5):791–8. doi: 10.1007/BF00028874.

32. Deng Z, Pang Y, Kong W, Chen Z, Wang X, Liu X, et al. A novel ABA-dependent dehydrin ERD10 gene from Brassica napus. DNA Sequence. 2005;16(1):28–35. doi: 10.1080/10425170500040180.

33. Alsheikh MK, Svensson JT, Randall SK. Phosphorylation regulated ion-binding is a property shared by the acidic subclass dehydrins. Plant Cell Envir. 2005;28(9):1114–22.

34. Kim SY, Nam KH. Physiological roles of ERD10 in abiotic stresses and seed germination of Arabidopsis. Plant Cell Rep. 2010;29(2):203–9. doi: 10.1007/s00299-009-0813-0.

35. Puhakainen T, Hess MW, Mäkelä P, Svensson J, Heino P, Palva ET. Overexpression of Multiple Dehydrin Genes Enhances Tolerance to Freezing Stress in Arabidopsis. Plant Mol Biol. 2004;54(5):743–53. doi: 10.1023/B:PLAN.0000040903.66496.a4.

36. Xie C, Zhang R, Qu Y, Miao Z, Zhang Y, Shen X, et al. Overexpression of *MtCAS31* enhances drought tolerance in transgenic Arabidopsis by reducing stomatal density. New Phytol. 2012;195(1):124–35. doi: 10.1111/j.1469-8137.2012.04136.x.

37. Guo X, Zhang L, Wang X, Zhang M, Xi Y, Wang A, et al. Overexpression of Saussurea involucrata dehydrin gene SiDHN promotes cold and drought tolerance in transgenic tomato plants. PLOS ONE. 2019;14(11):e0225090. doi: 10.1371/journal.pone.0225090.

38. Kumar M, Lee S-C, Kim J-Y, Kim S-J, Aye SS, Kim S-R. Over-expression of dehydrin gene, *OsDhn1*, improves drought and salt stress tolerance through scavenging of reactive oxygen species in rice (*Oryza sativa* L.). Journal of Plant Biology. 2014;57(6):383–93. doi: 10.1007/s12374-014-0487-1.

39. Šurbanovski N, Sargent DJ, Else MA, Simpson DW, Zhang H, Grant OM. Expression of *Fragaria vesca PIP* Aquaporins in Response to Drought Stress: *PIP* Down-Regulation Correlates with the Decline in Substrate Moisture Content. PLOS ONE. 2013;8(9):e74945. doi: 10.1371/journal.pone.0074945.

40. Jang JY, Kim DG, Kim YO, Kim JS, Kang H. An Expression Analysis of a Gene Family Encoding Plasma Membrane Aquaporins in Response to Abiotic Stresses in *Arabidopsis Thaliana*. Plant Mol Biol. 2004;54(5):713–25. doi: 10.1023/B:PLAN.0000040900.61345.a6.

41. He W-D, Gao J, Dou T-X, Shao X-H, Bi F-C, Sheng O, et al. Early Cold-Induced Peroxidases and Aquaporins Are Associated With High Cold Tolerance in Dajiao (*Musa* spp. ‘Dajiao’). Frontiers in Plant Science. 2018;9(282). doi: 10.3389/fpls.2018.00282.

42. Merlaen B, De Keyser E, Van Labeke M-C. Identification and substrate prediction of new *Fragaria x ananassa* aquaporins and expression in different tissues and during strawberry fruit development. Horticulture Research. 2018;5(1):20. doi: 10.1038/s41438-018-0019-0.

43. Tran L-SP, Nishiyama R, Yamaguchi-Shinozaki K, Shinozaki K. Potential utilization of NAC transcription factors to enhance abiotic stress tolerance in plants by biotechnological approach. GM Crops. 2010;1(1):32–9. doi: 10.4161/gmcr.1.1.10569.

44. Hao Y-J, Wei W, Song Q-X, Chen H-W, Zhang Y-Q, Wang F, et al. Soybean NAC transcription factors promote abiotic stress tolerance and lateral root formation in transgenic plants. The Plant Journal. 2011;68(2):302–13. doi: 10.1111/j.1365-313X.2011.04687.x.

45. An J-P, Li R, Qu F-J, You C-X, Wang X-F, Hao Y-J. An apple NAC transcription factor negatively regulates cold tolerance via CBF-dependent pathway. J Plant Physiol. 2018;221:74–80. doi: https://doi.org/10.1016/j.jplph.2017.12.009.

46. Zhuo X, Zheng T, Zhang Z, Zhang Y, Jiang L, Ahmad S, et al. Genome-Wide Analysis of the NAC Transcription Factor Gene Family Reveals Differential Expression Patterns and Cold-Stress Responses in the Woody Plant *Prunus mume*. Genes. 2018;9(10):494. Epub 2018/10/17. doi: 10.3390/genes9100494. PubMed PMID: 30322087; PubMed Central PMCID: PMCPMC6209978.

47. Sakuraba Y, Kim Y-S, Han S-H, Lee B-D, Paek N-C. The Arabidopsis Transcription Factor NAC016 Promotes Drought Stress Responses by Repressing *AREB1* Transcription through a Trifurcate Feed-Forward Regulatory Loop Involving NAP. The Plant Cell. 2015;27(6):1771–87. doi: 10.1105/tpc.15.00222.

48. Ohbayashi I, Lin C-Y, Shinohara N, Matsumura Y, Machida Y, Horiguchi G, et al. Evidence for a Role of ANAC082 as a Ribosomal Stress Response Mediator Leading to Growth Defects and Developmental Alterations in Arabidopsis. The Plant Cell. 2017;29(10):2644–60. doi: 10.1105/tpc.17.00255.

49. Zhou B, Zhang L, Ullah A, Jin X, Yang X, Zhang X. Identification of Multiple Stress Responsive Genes by Sequencing a Normalized cDNA Library from Sea-Land Cotton (*Gossypium barbadense* L.). PLOS ONE. 2016;11(3):e0152927. doi: 10.1371/journal.pone.0152927.

50. Apel K, Hirt H. REACTIVE OXYGEN SPECIES: Metabolism, Oxidative Stress, and Signal Transduction. Annu Rev Plant Biol. 2004;55(1):373–99. doi: 10.1146/annurev.arplant.55.031903.141701. PubMed PMID: 15377225.

51. Mittler R. Oxidative stress, antioxidants and stress tolerance. Trends Plant Sci. 2002;7(9):405–10. doi: https://doi.org/10.1016/S1360-1385(02)02312-9.

52. Sanmartin M, Drogoudi PD, Lyons T, Pateraki I, Barnes J, Kanellis AK. Over-expression of ascorbate oxidase in the apoplast of transgenic tobacco results in altered ascorbate and glutathione redox states and increased sensitivity to ozone. Planta. 2003;216(6):918–28. doi: 10.1007/s00425-002-0944-9.

53. Yamamoto A, Bhuiyan MNH, Waditee R, Tanaka Y, Esaka M, Oba K, et al. Suppressed expression of the apoplastic ascorbate oxidase gene increases salt tolerance in tobacco and *Arabidopsis* plants. J Exp Bot. 2005;56(417):1785–96. doi: 10.1093/jxb/eri167.

54. Fotopoulos V, Sanmartin M, Kanellis AK. Effect of ascorbate oxidase over-expression on ascorbate recycling gene expression in response to agents imposing oxidative stress. J Exp Bot. 2006;57(14):3933–43. doi: 10.1093/jxb/erl147.

55. Kim SJ, Ryu MY, Kim WT. Suppression of Arabidopsis RING-DUF1117 E3 ubiquitin ligases, AtRDUF1 and AtRDUF2, reduces tolerance to ABA-mediated drought stress. Biochem Biophys Res Commun. 2012;420(1):141–7. doi: https://doi.org/10.1016/j.bbrc.2012.02.131.

56. Kieber JJ, Rothenburg M, Roman G, Feldmann KA, Ecker JR. CTR1. a negtive regulator of the ethylene response pathway in arabidopsis, encodes a member of the Raf family of protein kinases. Cell. 1993;72(3):427–41.

57. Gao Z, Chen Y-F, Randlett MD, Zhao X-C, Findell JL, Kieber JJ, et al. Localization of the Raf-like Kinase CTR1 to the Endoplasmatic Reticulum of Arabidopsis through Participation in Ethylene Receptor Signalling Complexes. J Biol Chem. 2003;278:34725–32. doi: 10.1074/jbc.M305548200.

58. Chao Q, Rothenberg M, Solano R, Roman G, Terzaghi W, Ecker† JR. Activation of the Ethylene Gas Response Pathway in Arabidopsis by the Nuclear Protein ETHYLENE-INSENSITIVE3 and Related Proteins. Cell. 1997;89(7):1133–44. doi: https://doi.org/10.1016/S0092-8674(00)80300-1.

59. Shi Y, Tian S, Hou L, Huang X, Zhang X, Guo H, et al. Ethylene Signaling Negatively Regulates Freezing Tolerance by Repressing Expression of *CBF* and Type-A*ARR* Genes in *Arabidopsis*. The Plant Cell. 2012;24:2578–95.

60. Abuqamar S, Ajeb S, Sham A, Enan MR, Iratni R. A mutation in the expansin-like A2 gene enhances resistance to necrotrophic fungi and hypersensitivity to abiotic stress in Arabidopsis thaliana. Mol Plant Pathol. 2013;14(8):813–27. doi: https://doi.org/10.1111/mpp.12049.

61. Cosgrove DJ. Loosening of plant cell walls by expansins. Nature. 2000;407(6802):321–6. doi: 10.1038/35030000.

62. Chen T, Chen J-H, Zhang W, Yang G, Yu L-J, Li D-M, et al. BYPASS1-LIKE, A DUF793 Family Protein, Participates in Freezing Tolerance via the CBF Pathway in Arabidopsis. Frontiers in Plant Science. 2019;10(807). doi: 10.3389/fpls.2019.00807.

63. Le Hir R, Spinner L, Klemens Patrick AW, Chakraborti D, de Marco F, Vilaine F, et al. Disruption of the Sugar Transporters *AtSWEET11* and *AtSWEET12* Affects Vascular Development and Freezing Tolerance in *Arabidopsis*. Molecular Plant. 2015;8(11):1687–90. doi: 10.1016/j.molp.2015.08.007.

64. Essers L, Adolphs RH, Kunze R. A Highly Conserved Domain of the Maize *Activator* Transposase Is Involved in Dimerization. The Plant Cell. 2000;12:211–23.

65. Joly-Lopez Z, Forczek E, Vello E, Hoen DR, Tomita A, Bureau TE. Abiotic Stress Phenotypes Are Associated with Conserved Genes Derived from Transposable Elements. Frontiers in Plant Science. 2017;8(2027). doi: 10.3389/fpls.2017.02027.

66. Garcia ME, Lynch T, Peeters J, Snowden C, Finkelstein R. A small plant-specific protein family of ABI five binding proteins (AFPs) regulates stress response in germinating Arabidopsis seeds and seedlings. Plant Mol Biol. 2008;67(6):643–58. doi: 10.1007/s11103-008-9344-2.

67. Leung J, Giraudat J. ABSCISIC ACID SIGNAL TRANSDUCTION. Annu Rev Plant Physiol Plant Mol Biol. 1998;49(1):199–222. doi: 10.1146/annurev.arplant.49.1.199. PubMed PMID: 15012233.

68. Yu Y, Lv Y, Shi Y, Li T, Chen Y, Zhao D, et al. The role of phyto-melatonin and related metabolites in response to stress. Molecules. 2018;23(8):1887. doi: https://doi.org/10.3390/molecules23081887.

69. Gao H, Lu Z, Yang Y, Wang D, Yang T, Cao M, et al. Melatonin treatment reduces chilling injury in peach fruit through its regulation of membrane fatty acid contents and phenolic metabolism. Food Chem. 2018;245:659–66. doi: https://doi.org/10.1016/j.foodchem.2017.10.008.

70. Bajwa VS, Shukla MR, Sherif SM, Murch SJ, Saxena PK. Role of melatonin in alleviating cold stress in Arabidopsis thaliana. J Pineal Res. 2013;56(3):238–45. doi: https://doi.org/10.1111/jpi.12115.

